# Insights into the dynamic control of breathing revealed through cell-type-specific responses to substance P

**DOI:** 10.1101/754879

**Authors:** Nathan A Baertsch, Jan-Marino Ramirez

## Abstract

The rhythm generating network for breathing must continuously adjust to changing metabolic and behavioral demands. Here, we examine network-based mechanisms in the mouse preBӧtzinger complex using substance P, a potent excitatory modulator of breathing frequency and stability, as a tool to dissect network properties that underlie dynamic breathing. We find that substance P does not alter the balance of excitation and inhibition during breaths or the duration of the resulting refractory period. Instead, mechanisms of recurrent excitation between breaths are enhanced such that the rate that excitation percolates through the network is increased. Based on our results, we propose a conceptual framework in which three distinct phases, the inspiratory phase, refractory phase, and percolation phase, can be differentially modulated to influence breathing dynamics and stability. Unravelling mechanisms that support this dynamic control may improve our understanding of nervous system disorders that destabilize breathing, many of which are associated with changes in brainstem neuromodulatory systems.

## Introduction

Rhythmicity is ubiquitous in the brain, important for many high order functions such as consciousness, attention, perception, and memory (Basar and Duzgun, 2016, Basar and Guntekin, 2008, Colgin, 2016, Hanslmayr et al., 2016, Kiehn, 2016, Neske, 2015, Palva and Palva, 2018, Paton and Buonomano, 2018), as well as vital rhythmic motor behaviors including chewing, locomotion, and breathing (Grillner and El Manira, 2015, Kiehn, 2016, Narayanan and DiLeone, 2017, Ramirez and Baertsch, 2018a, Wyart, 2018, Nakamura et al., 2004). Rhythms generated by the brain are diverse, as are the underlying rhythm generating network- and cellular-level mechanisms (Paton and Buonomano, 2018). Indeed, rhythmicity can occur on time scales ranging from milliseconds to days (Golombek et al., 2014). However, most neuronal rhythms must be flexible, able to increase or decrease the rate of oscillation or the degree of synchronization to match changes in physiological or cognitive demands (Brittain et al., 2014, Ramirez and Baertsch, 2018b). Thus, understanding principles that allow dynamic control of rhythm generating networks may provide important insights into the regulation of diverse brain functions.

For half a century, investigations of the networks and cellular mechanisms that generate breathing have provided valuable, generalizable insights into the origins and control of neural rhythmicity (Del Negro et al., 2018, Feldman and Kam, 2015, Ramirez and Baertsch, 2018b, Wyman, 1977, Cohen, 1981, Ezure, 1990, Long and Duffin, 1986, Milsom, 1991). Inspiration, the dominant phase of breathing in mammals (Jenkin and Milsom, 2014), is generated by a spatially dynamic network located bilaterally along the ventrolateral medulla (Baertsch et al., 2019). A region that is both necessary and sufficient for inspiration, the pre-Bötzinger Complex (preBötC), is autorhythmic and forms the core of this network (Smith et al., 1991, Tan et al., 2008, Vann et al., 2018). Like many rhythmic networks, a critical characteristic of the preBӧtC is that the frequency of its output is dynamic - the rate of breathing changes during e.g. sleep/wake states, exercise, environmental challenges, and orofacial behaviors such as feeding and vocalization (Moore et al., 2013, Ramirez et al., 2016). Excitatory and inhibitory inputs from other brain regions, as well as neuromodulation, can potently facilitate or depress the frequency of breathing (Doi and Ramirez, 2008, Zuperku et al., 2017). Yet, elucidating how these influences alter the network- and cellular-level rhythm generating mechanisms within the preBӧtC remains a challenge (Dick et al., 2018).

Glutamatergic synaptic interactions among preBötC interneurons allow this sparsely connected network (Carroll and Ramirez, 2013, Schwab et al., 2010) to periodically synchronize and are therefore obligatory for rhythmogenesis (Ge and Feldman, 1998). However, if left unrestrained, feed-forward excitation in the network leads to hyper synchronization during inspiratory bursts, which subsequently causes a prolonged period of reduced network excitability (Baertsch et al., 2018, Kottick and Del Negro, 2015). This refractory phase delays the onset of the next inspiratory burst and, as a result, the frequency of breathing becomes very slow. Inhibitory interactions within the preBӧtC are critical for limiting synchronization during bursts (Harris et al., 2017) and reducing the subsequent refractoriness of the network (Baertsch et al., 2018). Indeed, roughly 40% of inspiratory preBötC neurons are inhibitory (GABAergic and/or glycinergic) (Oke et al., 2018, Winter et al., 2009) and by regulating the excitability of glutamatergic neurons during inspiratory bursts, these neurons play an important role in controlling breathing frequency.

However, controlling the inspiratory burst itself is not the only mechanism that regulates the inspiratory rhythm. During the time between bursts, referred to as the inter-burst interval (IBI), recurrent excitatory synaptic connections within the preBӧtC are thought to give rise to a gradual increase in network excitability that drives the onset of the next burst (Del Negro et al., 2018). In this model, spontaneous spiking activity in a small subset of excitatory preBӧtC neurons begins to percolate stochastically through the network, gradually recruiting more spiking activity among interconnected excitatory neurons (Kam et al., 2013b). During this percolation phase, activation of membrane voltage- and calcium-dependent conductances in an increasing number of neurons causes the excitation to become exponential, culminating in an inspiratory burst (Del Negro et al., 2010, Ramirez et al., 2016). The gradual increase in spiking activity during this phase, or “pre-inspiratory ramp”, is thought to be primarily mediated by a subset of glutamatergic preBӧtC interneurons that have enhanced excitability. Derived from V0-lineage precursors, these neurons express the transcription factor developing brain homeobox 1 protein (Dbx1) during development (referred to here as “Dbx1 neurons”) (Bouvier et al., 2010, Gray et al., 2010, Picardo et al., 2013, Wu et al., 2017). How this process of recurrent excitation may contribute to the dynamic regulation of inspiratory frequency is not well understood.

Here, we examine network- and cellular-level changes in the inspiratory rhythm generator that underlie dynamic frequency responses to the excitatory neuromodulator substance P (SP). A member of the tachykinin neuropeptide family, SP is a key mediator of many physiological and neurobiological processes (e.g. Mantyh, 2002). For breathing, SP regulates the stability of the respiratory rhythm (Ben-Mabrouk and Tryba, 2010, Yeh et al., 2017) as well as respiratory responses to hypoxia (Chen et al., 1990, Ptak et al., 2002). The endogenous receptor for SP, neurokinin 1 receptor (NK_1_R), is expressed on only ∼5-7% of CNS neurons (Mantyh, 2002), but is enriched in the preBӧtC (Gray et al., 1999, Schwarzacher et al., 2011). SP binding to NK_1_R causes excitation of preBӧtC neurons through coupling with voltage-independent cation channels (Hayes and Del Negro, 2007, Ptak et al., 2009), including *sodium leak channel, non-selective (Nalcn)*. Disruption of this ion channel causes pathological respiratory instability (Yeh et al., 2017). SP also promotes robust facilitation of inspiratory frequency (Gray et al., 1999). Therefore, SP is an ideal tool to explore how changes in network interactions during the inspiratory cycle influence breathing stability and promote dynamic regulation of breathing frequency.

By combining electrophysiological, optogenetic, and pharmacological techniques, we find that SP differentially influences the refractory and recurrent excitation phases of the inspiratory rhythm through cell-type-specific effects. We conclude that phase-specific and differential modulation of excitatory and inhibitory network interactions is a key mechanism that allows the frequency of this vital rhythmogenic network to be dynamically controlled.

## Results

### SP has differential effects on the refractory and percolation phases of the preBӧtC rhythm

To explore how the neuromodulator SP increases inspiratory frequency at the network level, integrated preBӧtC population activity was recorded in horizontal brainstem slices (Anderson et al., 2016, Baertsch et al., 2019) from Dbx1^ERT2Cre^;Rosa26^ChR2EYFP^ neonatal mice during bath application of 0.5-1.0µM SP (n=6). As expected (Pena and Ramirez, 2004), inspiratory burst frequency increased from 0.23±0.02Hz to 0.31±0.03Hz (p=0.003) during steady state SP (>∼3min post bath application). To determine whether the refractory period is modulated by SP, brief light pulses (200ms, 0.5mW/mm^2^) were delivered randomly during the inspiratory cycle at baseline and in SP. In each condition, the probability of light-evoking a burst in the contralateral preBӧtC was quantified as a function of elapsed time from the preceding spontaneous population burst and compared to the cumulative distribution of spontaneous inter-burst intervals (IBIs). A representative experiment is shown in Fig. 1A and B, and the average data are shown in Fig. 1C. Evoked bursts were rare if a light pulse occurred immediately following a spontaneous population burst. However, the probability of a light-evoked burst increased with elapsed time until ∼2 sec following a spontaneous burst when bursts could be evoked with nearly every light pulse (Fig. 1A,B). The end of this ∼2 sec refractory period coincided with a large increase in the number of spontaneous IBIs (Fig.1B,C), indicating that this period of reduced preBӧtC excitability precludes both light-evoked and spontaneous preBӧtC burst generation, thereby preventing very short IBIs and fast inspiratory rhythms (Baertsch et al., 2018). However, despite a frequency increase of 31.9±5.6%, the refractory period was not altered by SP (non-linear regression analysis; p>0.05). In SP, IBIs remained limited by the refractory period, but spontaneous bursts occurred more quickly and more consistently following the end of the refractory period. As a result, the average IBI became shorter (4.5±0.4sec to 3.5±0.5sec; paired t-test; p<0.0002) and less variable (SD of 1.3±0.1 to 0.84±0.1; p<0.0139) in SP (Fig. 1C). Together these data suggest that SP increases the frequency and regularity of the inspiratory rhythm through differential modulation of two inspiratory phases: The refractory phase remains unchanged, while the duration of the recurrent excitation or percolation phase, which promotes the onset of the subsequent burst, is reduced.

**Figure 1:**
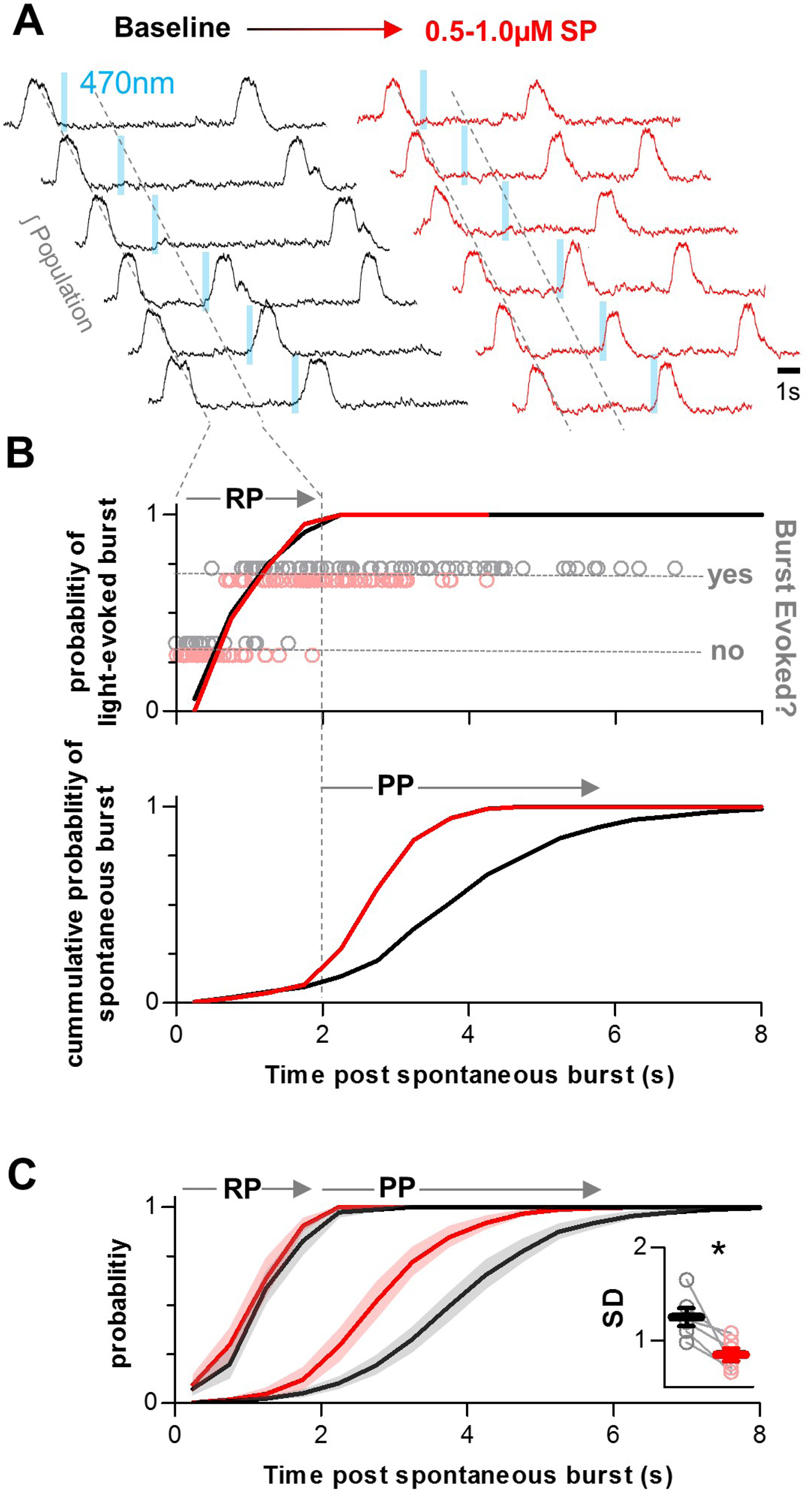
Differential modulation of the refractory phase (RP) and recurrent excitation or percolation phase (PP) of the inspiratory rhythm by SP. **A**) Representative integrated preBotC population recordings from a Dbx1^ERT2Cre^;Rosa26^ChR2EYFP^ horizontal slice during photostimulation of Dbx1 neurons under baseline conditions (black) and in SP (red). **B**) Quantified data from the experiment in A showing time-dependent changes in the probabilities of evoking a burst (top) and of a spontaneous burst occurring (bottom). **C**) Group data from n=6 experiments. Spontaneous and evoked probability curves were compared under baseline conditions and in SP using non-linear regression analysis. Inset shows the standard deviation (SD) of the inter-burst intervals under baseline conditions and in SP (paired, two tailed t-test).

### Inspiratory spiking patterns of excitatory and inhibitory neurons in the preBӧtC

Next, we explored the spiking patterns of individual excitatory and inhibitory preBӧtC neurons to identify mechanisms that may underlie differential modulation of the refractory and percolation phases in the preBӧtC network. Inspiratory spiking activity was recorded from n=29 neurons located in the preBӧtC. Horizontal slices from Vglut2^Cre^;Rosa26^ChR2EYFP^ and Vgat^Cre^;Rosa26^ChR2EYFP^ mice were used so that recorded neurons could be identified as excitatory or inhibitory based on depolarizing responses to light (Baertsch et al., 2018, Baertsch et al., 2019). Recorded neurons were also fluorescently labelled using patch pipets containing AlexaFluor568 to mark their anatomical locations (Fig. 2A). There was considerable variability among inspiratory neurons (maximal spiking activity during inspiration), with respect to spike frequency, burst duration, and burst shape; and for any given neuron there was considerable burst-to-burst stochasticity (Carroll and Ramirez, 2013, Carroll et al., 2013). However, excitatory neurons could be clearly grouped based on the presence or absence of spiking during the IBI that typically increases in frequency, or “ramps”, before the subsequent inspiratory burst – often referred to as “pre-inspiratory (pre-I)” activity. In contrast, we did not identify any inspiratory inhibitory neurons in the preBӧtC with pre-I spiking (Fig. 2B). Excitatory neurons with pre-I spiking (n=9) had inspiratory spike frequencies ranging from 12.0 to 50.8Hz (mean: 28.0±4.4Hz) and burst durations ranging from 272 to 763ms (mean: 467±48ms). Similarly, excitatory neurons without pre-I spiking (n=13) had inspiratory spike frequencies ranging from 11.3 to 58.2Hz (mean: 34.9±4.3Hz) and burst durations ranging from 242 to 698ms (mean: 419±34ms). Inhibitory neurons (n=7) also had considerable variability in spike frequency (16.4 to 82.9Hz; mean: 48.6±8.6Hz) and burst duration (144 to 517ms; mean: 316±54ms) (Fig. 2C,D).

**Figure 2:**
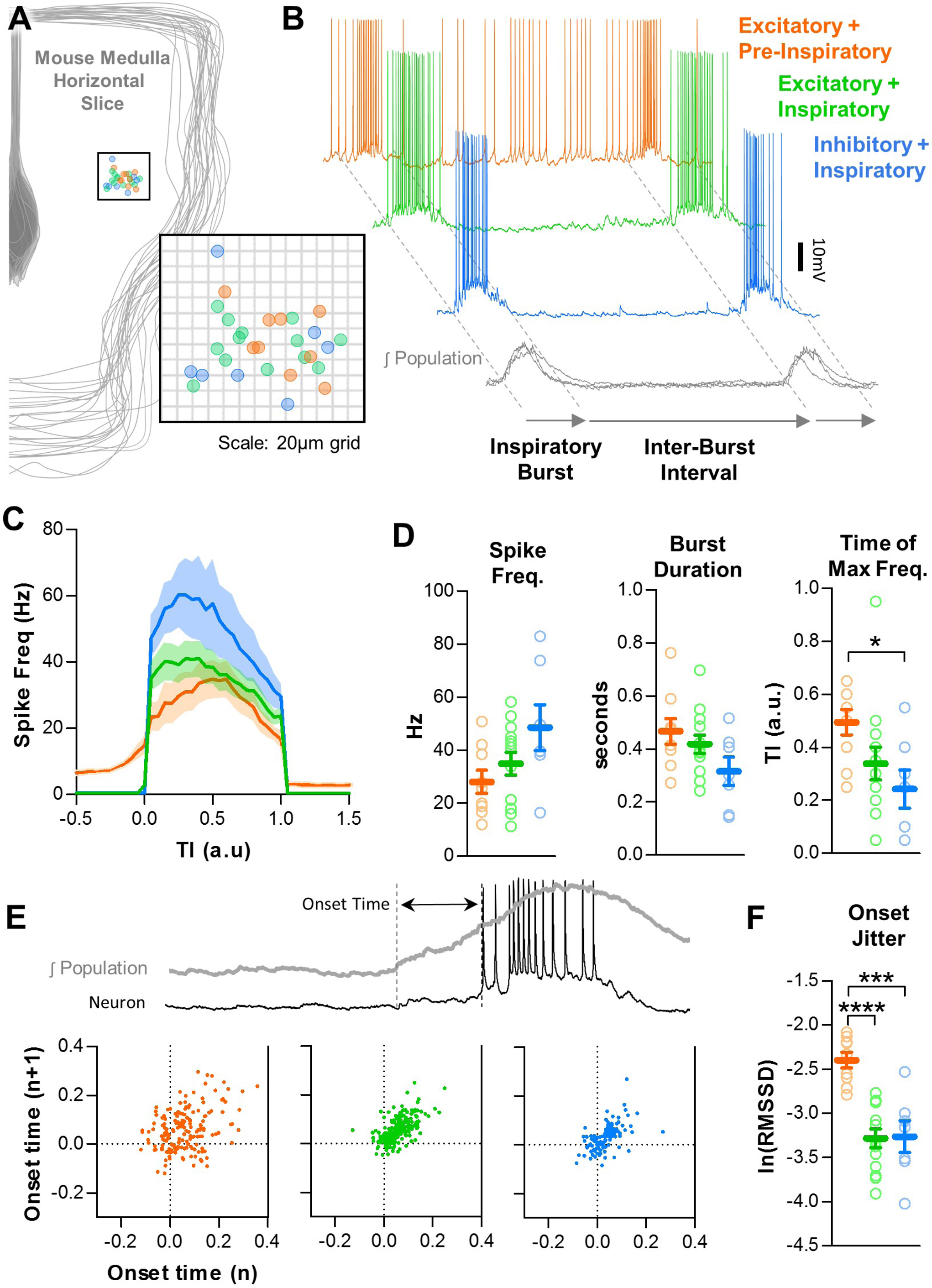
Baseline spiking patterns of excitatory and inhibitory neurons in the preBotC. **A**) Anatomical locations of n=29 recorded preBotC neurons. **B**) Example traces of an excitatory neuron with pre-I spiking (orange) and without pre-I spiking (green) and an inhibitory neuron (blue) during the inspiratory burst and inter-burst interval (IBI). **C)** Quantified spike frequency as a function of time (normalized to burst duration) from n=9 pre-I excitatory, n=13 non pre-I excitatory, and n=7 inhibitory neurons. **D**) Quantified mean spike frequency, duration, and shape of inspiratory bursts generated by each type of neuron (one-way ANOVA with Bonferroni poc hoc tests). **E**) Example quantification of neuronal burst onset time relative to the preBӧtC population and Poincaré plots showing onset time variability from n=9 pre-I excitatory, n=13 non pre-I excitatory, and n=7 inhibitory neurons (20 inspiratory bursts/neuron). **F**) Burst onset time variability or “jitter” quantified as the natural log of the root mean square of successive differences (one-way ANOVA with Bonferroni post hoc tests).

Inspiratory neuron types also differed in burst shape (Fig. 2C,D) and burst onset variability (Fig. 2E,F). Inhibitory neurons had a much more pronounced decrementing spiking pattern than excitatory neurons with maximal spike frequencies occurring at 24±7% of the inspiratory burst duration. Excitatory neurons with pre-I spiking exhibited a rounded burst pattern with maximal spike frequencies occurring at 49±5% of the inspiratory burst duration, whereas excitatory neurons without pre-I spiking were slightly more decrementing with maximal spike frequencies occurring at 34±6% of the inspiratory burst duration. We also quantified the time between the beginning of each preBӧtC population burst and the onset of the corresponding neuronal burst (i.e. “onset time”). Poincaré plots of onset times for each inspiratory neuron type are shown in Fig. 2E and burst onset variability (quantified as the natural log of the root mean square of successive differences, ln(RMSSD)) is shown in Fig. 2F. Overall, average onset times did not differ among neuron types (p>0.05); however excitatory neurons with pre-I spiking had more cycle-to-cycle variability in burst onset times than excitatory neurons without pre-I spiking or inhibitory neurons (p<0.001).

### Effects of SP on excitatory pre-inspiratory neurons in the preBӧtC are phase-dependent

Spiking activity of pre-I neurons is expected to contribute to both the refractory and percolation phases of the inspiratory rhythm. During inspiratory bursts, spiking of these neurons contributes to synchronization, which promotes the subsequent refractoriness of the network (Baertsch et al., 2018). Following the refractory phase, it is thought that pre-I spiking of these neurons during the IBI facilitates positive-feedback recurrent excitation in the network, which builds up until another inspiratory burst in generated and the cycle restarts (Del Negro et al., 2018). Thus, changes in spiking during inspiratory bursts are predicted to alter the duration of the subsequent IBI through modulation of the refractory phase, whereas changes in pre-inspiratory spiking are predicted to alter the IBI by changing the rate of feed-forward excitation during the percolation phase. We examined SP-induced changes in the spiking activity of pre-I neurons during inspiratory bursts and during the IBI. A representative recording is shown in Fig. 3A. Unexpectedly, SP had very little effect on spiking during inspiratory bursts (Fig. 3B). Changes in burst spike frequency (28.0±4.4Hz to 29.6±3.8Hz, p>0.05) and burst duration (467±78ms to 488±45ms; p>0.05) were small and inconsistent (Fig. 3B,C). In contrast, during the IBI, SP increased the average spiking frequency of pre-I neurons from 5.1±1.0Hz to 9.9±1.8Hz (p<0.01), and in all cases increased the slope of the pre-inspiratory ramp (average of 1.7±0.8 to 3.7±1.9Hz/sec), although this did not reach statistical significance (p=0.113) due to the large variability among neurons (Fig. 3D,E). Changes in IBI spike frequency and slope induced by SP were coincident with a significant decrease in the duration of the IBI (Fig. 3E). Thus, these phase-dependent changes in spiking activity at the level of individual pre-I excitatory neurons likely contribute to the differential effects of SP on the refractory and percolation phases observed at the network level.

**Figure 3:**
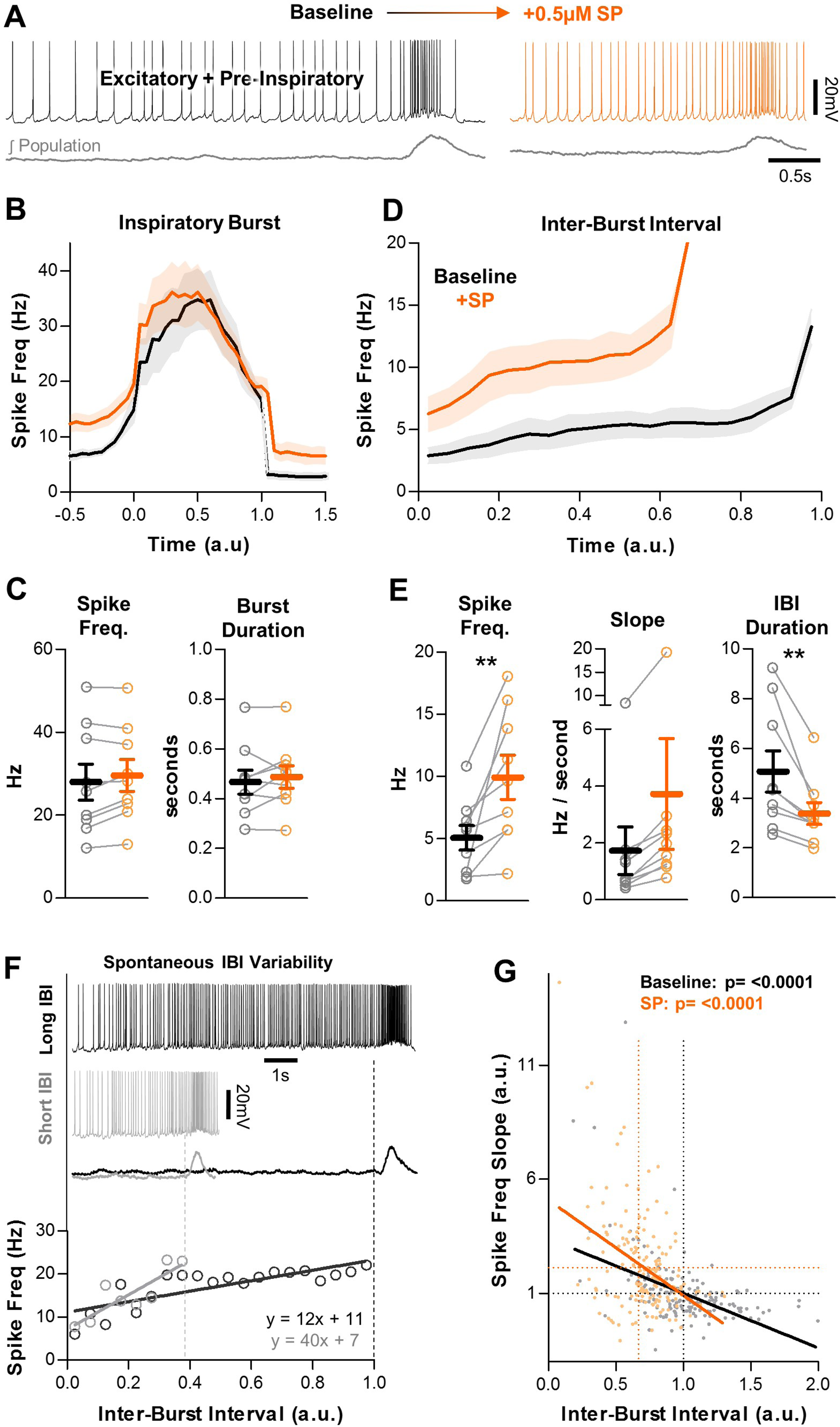
Effects of SP on pre-I excitatory neurons in the preBӧtC. **A**) Example intracellular recording from a pre-I neuron at baseline (black) and in SP (orange) with corresponding integrated preBotC population activity (grey). **B**) Quantified spike frequency as a function of time (normalized to inspiratory burst duration) in n=9 pre-I neurons. **C**) Mean spike frequency and burst duration of pre-I neurons (paired, two tailed t-tests). **D**) Quantified spike frequency vs. time (normalized to IBI duration) during the inter-burst interval showing changes in pre-inspiratory ramp activity induced by SP. **E**) Mean spike frequency, pre-inspiratory ramp slope, and IBI duration (paired, two tailed t-tests). **F**) Example spiking of a pre-I neuron during a long (black) and short (grey) inter-burst interval under baseline conditions (top), and quantified pre-inspiratory slope during each IBI (below) **G**) Inverse relationship between the slope of pre-inspiratory spiking and the length of the IBI from n=9 pre-I neurons (20 consecutive IBIs/neuron) at baseline and in SP (parameters normalized to baseline values) (linear regression analysis).

Since SP also reduced the variability of the IBI at the network level (see Fig. 1B), we examined the relationship between the duration of individual IBIs and the slope of pre-inspiratory spiking activity. An example recoding of an excitatory pre-inspiratory neuron during a long IBI (black) and a short IBI (grey), and the quantified spike frequency over the duration of each IBI, is shown in Fig. 3F. Group data for n=9 neurons is shown in Fig. 3G. 20 consecutive inspiratory cycles were analyzed for each neuron and the duration of each IBI was compared to the pre-inspiratory slope during that cycle. To highlight effects related to cycle-to-cycle variability, values were normalized to the average baseline IBI duration and pre-inspiratory slope for each neuron, respectively. We found that, under control conditions and in SP, there was a significant inverse relationship between the duration of a given IBI and the slope of the corresponding pre-inspiratory ramp, such that pre-inspiratory spiking activity at the level of individual neurons can predict the duration between inspiratory bursts at the network level.

### SP recruits a subpopulation of excitatory preBӧtC neurons to exhibit pre-inspiratory spiking

Unlike pre-I neurons, excitatory neurons that are silent during the IBI are unable to participate in the feed-forward process of recurrent excitation because they lack pre-inspiratory spiking activity. However, these neurons are expected to contribute to network synchronization during inspiratory bursts, and as a result they have the potential to modulate the refractory period. In response to SP, excitatory neurons that did not spike during the IBI under baseline conditions exhibited two distinct phenotypes. Some (8/13) remained silent during the IBI (teal), whereas others (5/13) developed pre-inspiratory spiking (green) (Fig. 4A). Excitatory neurons that were not recruited to spike during the IBI also had no change in spike frequency (38.6±5.0 to 37.1±4.9Hz; p>0.05) or burst duration (430±53 to 428±51ms, p>0.05) during inspiratory bursts (Fig. 4B,C), despite a shortened IBI duration (p<0.05). Since these neurons had no change in spiking throughout the inspiratory cycle, it is unlikely that they contribute to SP-induced frequency facilitation of the inspiratory rhythm. In neurons that were recruited to spike during the IBI, spiking frequency increased from 0 to 6.6±1.4Hz (p<0.01). In SP, these neurons exhibited a pre-inspiratory ramp (4.1±1.7 Hz/sec), which was coincident with a shorter IBI duration (p<0.05) (Fig. 4D,E). During individual inspiratory cycles, the slope of the SP-induced pre-inspiratory ramp had a significant inverse relationship with the duration of the IBI (Fig. 4F). During inspiratory bursts, spiking patterns did not change in spike frequency (28.9±7.8 to 31.9±6.6Hz, p>0.05), burst duration (399±28 to 441±36ms, p>0.05), or burst shape (Fig. 4B,C). Thus, a subpopulation of non pre-I excitatory neurons develops pre-I activity in SP without a change in spiking activity during bursts, suggesting that the number of neurons that can participate in the percolation phase increases in the presence of SP, without significant effects on the amount of excitation during bursts and the resulting refractory period.

**Figure 4:**
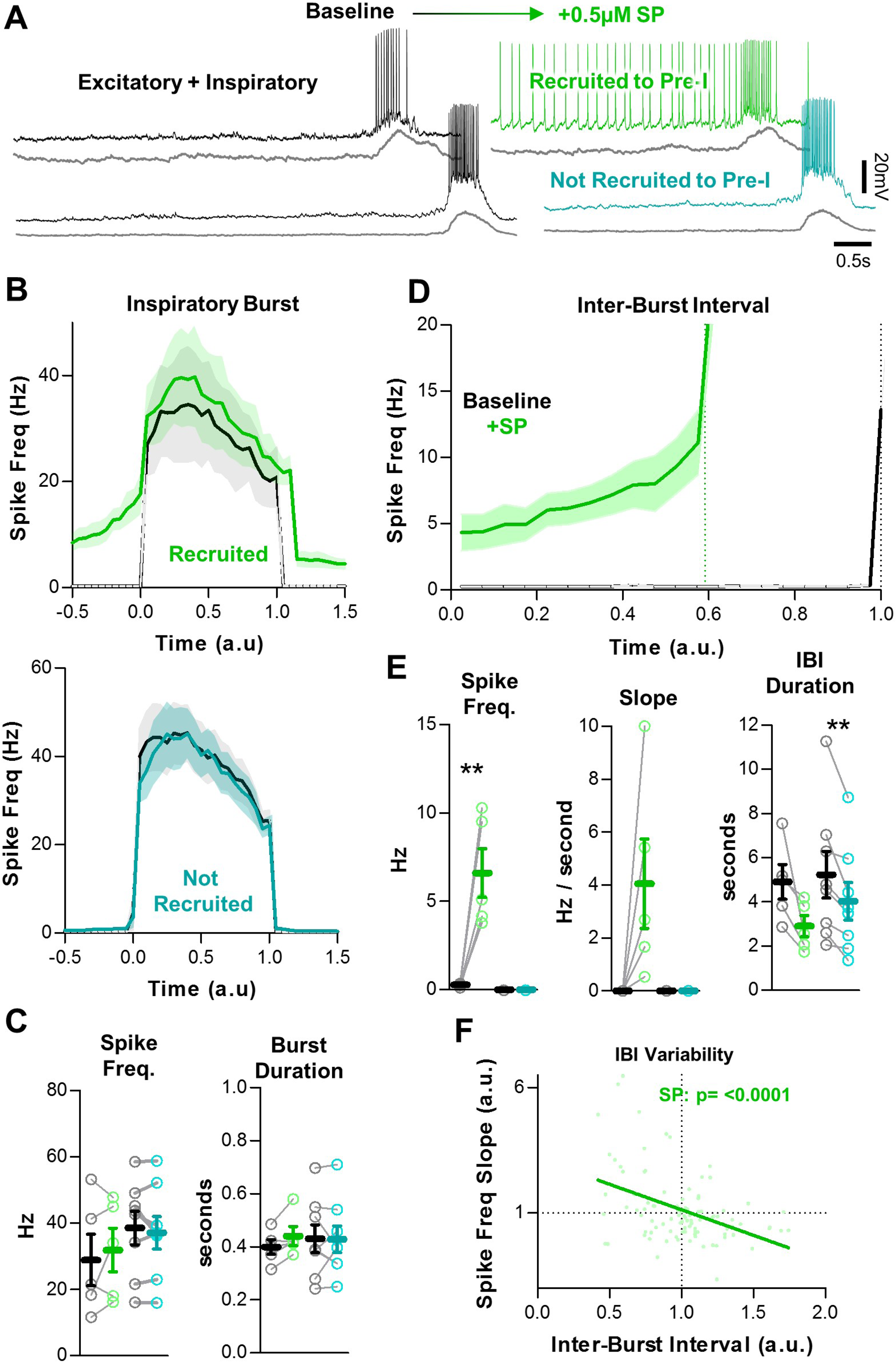
SP recruits a subpopulation of excitatory preBӧtC neurons to participate in the percolation phase. **A**) Example intracellular recordings from two excitatory neurons, one that develops pre-I activity in SP (green) and one that does not (teal). Corresponding integrated preBӧtC population activity is shown below each trace (grey). **B**) Quantified spike frequency as a function of time (normalized to baseline inspiratory burst duration) in n=5 excitatory neurons that were recruited to pre-I (top) and n=8 excitatory neurons that were not recruited to pre-I (bottom). **C**) Mean spike frequency and burst duration in both neuron groups (paired, two-tailed t-tests). **D**) Quantified spike frequency vs. time (normalized to baseline IBI duration) during the inter-burst interval showing the recruitment of pre-inspiratory ramp activity by SP. **E**) Mean spike frequency, pre-inspiratory ramp slope, and IBI duration in both neuron groups (paired, two tailed t-tests). **F**) Inverse relationship between the slope of pre-inspiratory spiking and the length of the IBI in SP from n=5 excitatory neurons that were recruited to pre-I (20 consecutive IBIs/neuron) (linear regression analysis).

### SP does not change spiking activity of inhibitory inspiratory neurons in the preBӧtC

In contrast to excitatory neurons, the activity of inhibitory neurons during inspiratory bursts reduces network synchronization and the refractory period (Baertsch et al., 2018). Since the RP of the preBӧtC network was not changed by SP (see Fig. 1), we hypothesized that inhibition during bursts would also be unchanged by SP. To test this, we recorded spiking activity from n=7 inhibitory preBӧtC neurons under baseline conditions and following application of SP. A representative recoding is shown in Fig. 5A. In the presence of SP, spike frequency and burst duration of inspiratory inhibitory neurons did not change during bursts (48.6±8.6 to 43.5±6.7Hz, p>0.05; and 316±54 to 328±56ms, p>0.05, respectively), despite a coincident shortening of the IBI (p<0.05) (Fig. 5B,C). Spiking activity of inhibitory neurons also did not change during the IBI, since all of the recorded neurons remained silent between inspiratory bursts. Thus, in response to SP, inhibitory neurons in the preBӧtC had no change in spiking throughout the inspiratory cycle and are therefore unlikely to play a role in SP-induced facilitation of inspiratory frequency.

**Figure 5:**
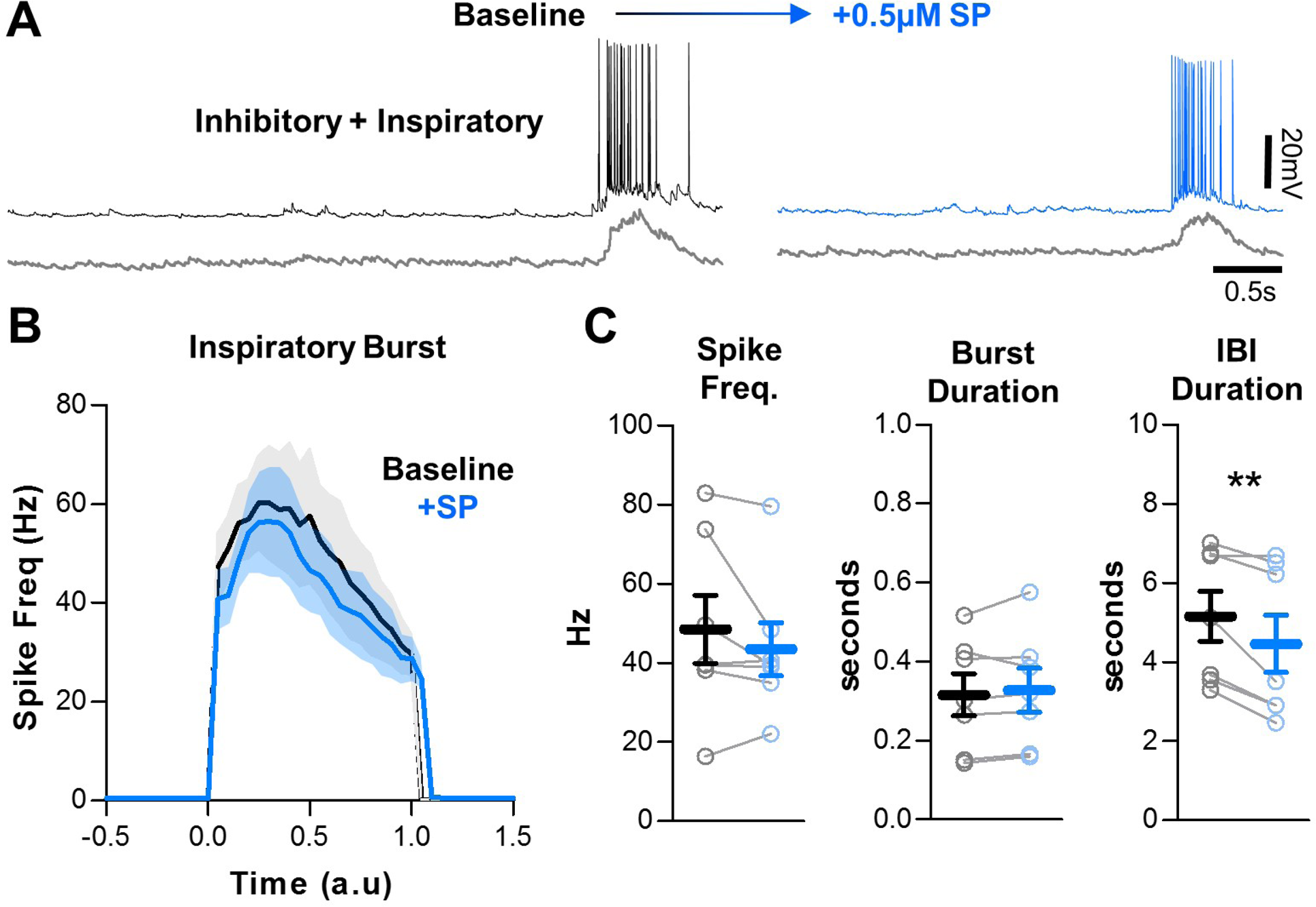
SP does not change inhibitory network interactions. **A**) Example intracellular recordings from an inhibitory preBӧtC neuron under baseline conditions (black) and in SP (blue) with corresponding integrated preBӧtC population activity is shown below (grey). **B**) Quantified spike frequency as a function of time (normalized to baseline inspiratory burst duration) in n=7 inhibitory neurons. **C**) Mean spike frequency, burst duration, and IBI (paired, two-tailed t-tests).

### SP increases stochasticity among excitatory preBӧtC neurons during inspiratory bursts

Next, we sought to unravel potential mechanisms that may prevent SP from causing hyper-synchronization of the preBӧtC network and increased refractory times. Since synchronization is often reduced with increased stochasticity (Carroll and Ramirez, 2013, Harris et al., 2017, Zerlaut and Destexhe, 2017), we compared the cycle-to-cycle variability in burst onset times (see Fig. 2E) under baseline conditions and in the presence of SP (Fig. 6). Although average burst onset times were not significantly altered by SP for any neuronal type (p>0.05), SP did have effects on burst onset stochasticity. This is demonstrated as greater dispersions in the Poincaré plots shown in Fig. 6A. However, these SP-induced changes in burst onset variability (i.e. “onset jitter”) differed across neuronal types. Burst onset jitter of excitatory pre-I neurons, which was generally high under baseline conditions (see Fig. 2F), did not change significantly in SP (p>0.05) (Fig.6B). In contrast, excitatory neurons that did not exhibit pre-I spiking and had relatively low onset jitter under baseline conditions exhibited increased onset jitter in the presence of SP (p<0.05). Among this group of excitatory neurons, burst onset jitter was increased by SP regardless of whether or not the neuron was recruited to develop pre-inspiratory spiking. Inhibitory preBӧtC neurons, on the other hand, had relatively inconsistent changes in burst onset variability induced by SP, with no change in mean onset jitter (p>0.05). These results suggest that SP increases the stochasticity of burst onset in a subgroup of preBӧtC excitatory neurons, which may help prevent this excitatory neuromodulator from causing hyper-synchronization among preBӧtC neurons during inspiratory bursts.

**Figure 6:**
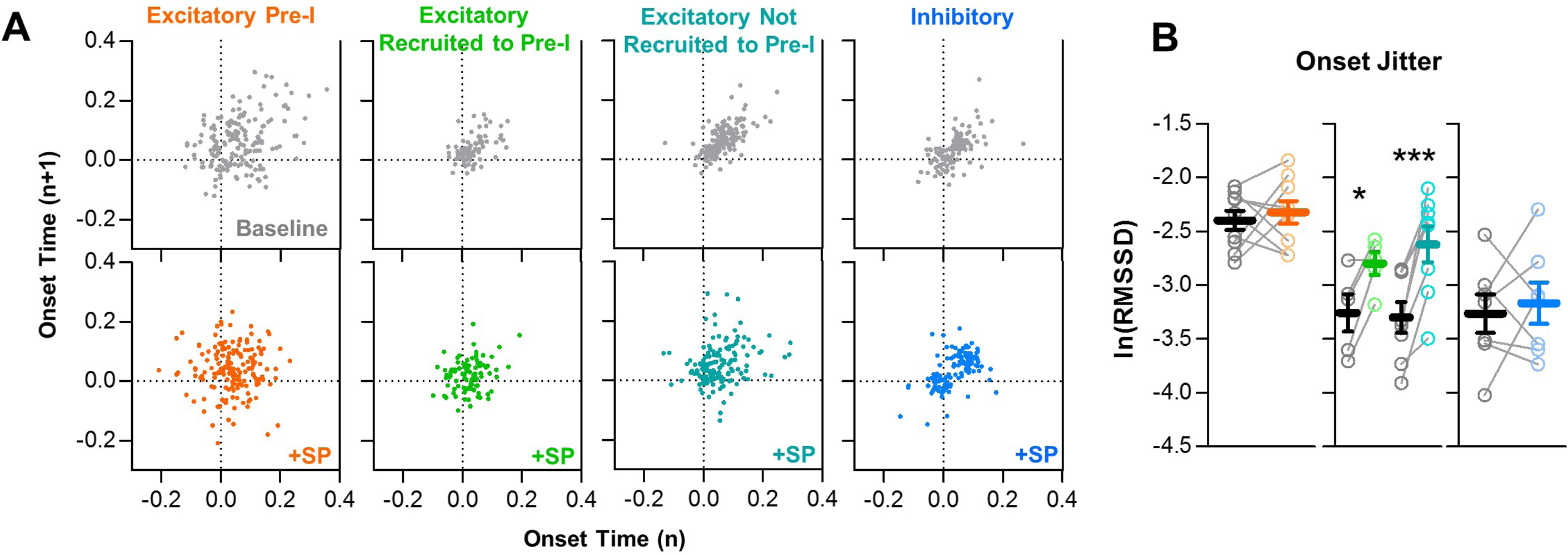
SP increases onset variability among non pre-I excitatory neurons during preBӧtC bursts. **A**) Poincaré plots of burst-to-burst variability in onset times under baseline conditions (grey) and in SP for n=9 excitatory, pre-I (orange); n=5 excitatory, recruited to pre-I (green); n=8 excitatory, not recruited to pre-I (teal); and n=7 inhibitory (blue) neurons. **B**) Burst onset time variability or “jitter”, quantified as the natural log of the root mean square of successive differences, at baseline and in SP for each neuron type. (paired, two tailed t-tests).

### Inspiratory neurons rostral to the preBӧtC have heterogeneous responses to SP

Neurons with inspiratory activity are not confined to the preBӧtC but are distributed along the ventral respiratory column (VRC) (Barnes et al., 2007, Zuperku et al., 2019). Indeed, the inspiratory network seems to be spatially dynamic since excitatory neurons located rostral to the preBӧtC can be conditionally recruited to participate in the inspiratory rhythm (Baertsch et al., 2019). This rostral expansion of the active inspiratory network is associated with an increase in the excitation/inhibition ratio, longer refractory times, and slower inspiratory frequencies (Baertsch et al., 2019). Therefore, we explored whether the opposite may also occur: Could the size of the active inspiratory network shrink, and could this be another mechanism that prevents SP from causing increased excitation during inspiratory bursts? To test this, we recoded spiking activity from n=16 inspiratory neurons located in the rostral VRC (n=5 excitatory, n=7 inhibitory, n=4 unknown). The anatomical locations of these neurons relative to the preBӧtC neurons described above are shown in Fig. 7A. Overall, spiking activity patterns and responses to SP were less consistent among rostral neurons than preBӧtC neurons. To convey this heterogeneity, spike rasters for each rostral neuron over 20 consecutive inspiratory bursts are shown in Fig. 7B,C,D. Despite this variability, spiking frequency during bursts was reduced by SP in all (5/5) rostral excitatory neurons (-55.6±13.5% change from 11.1±4.7 to 6.7±4.0Hz; p<0.05) (Fig. 8A,C), whereas changes were inconsistent among inhibitory rostral neurons with no change on average (39.5±34.7% change from 5.3±1.5 to 6.1±1.5Hz; p>0.05) (Fig. 8B,C). Thus, the potential contribution of rostral excitatory neurons to synchronization of the inspiratory rhythm was reduced by SP, while inhibitory influences were relatively unchanged. Burst onset variability of both excitatory and inhibitory inspiratory neurons rostral of the preBӧtC was generally high under baseline conditions, and it was further increased by SP (p<0.05) (Fig. 8D,E).

**Figure 7:**
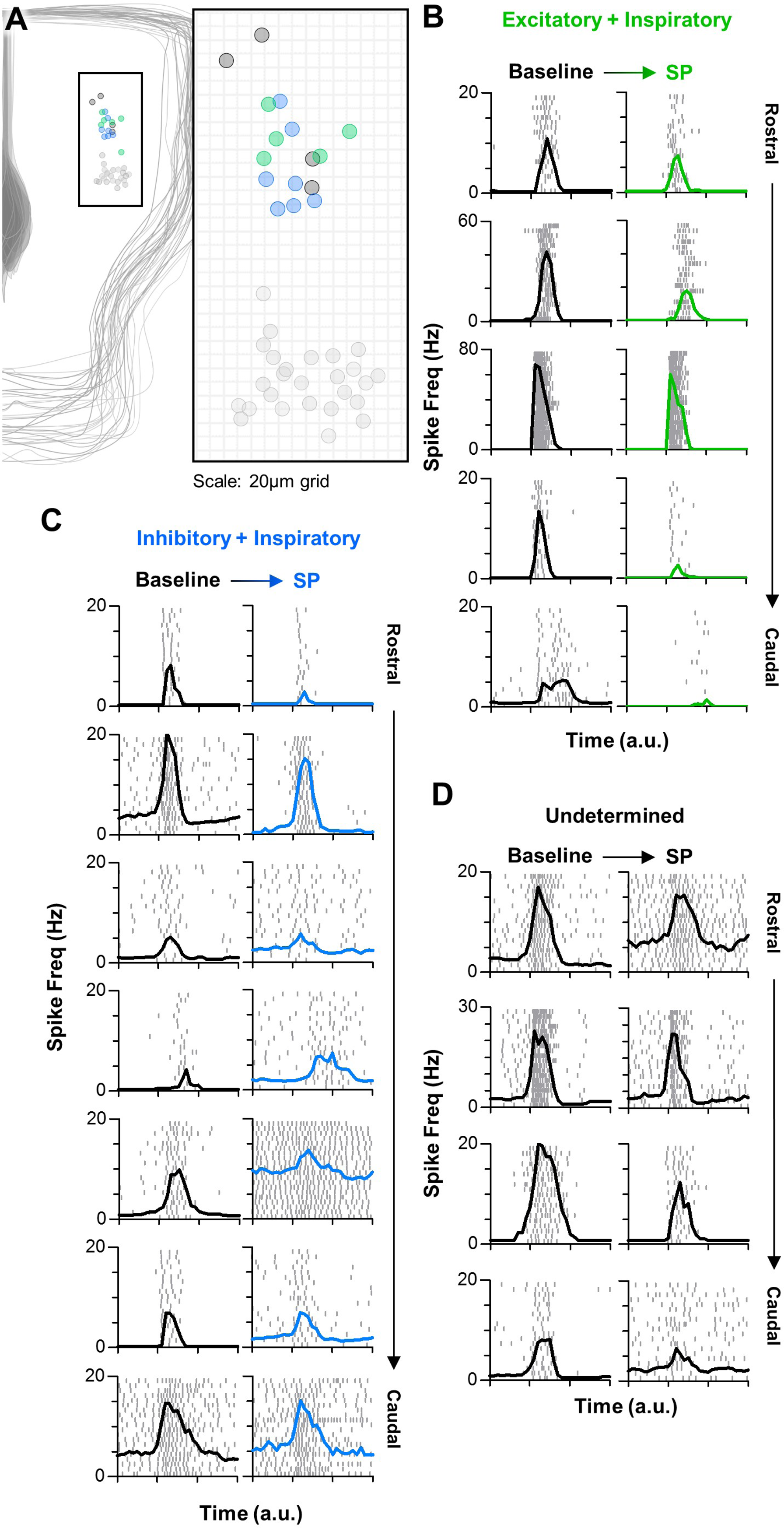
Inspiratory neurons rostral of the preBӧtC have varied responses to SP. **A**) Anatomical locations of n=16 rostral inspiratory neurons [n=5 excitatory (green), n=7 inhibitory (blue), and n=4 unknown (black)], relative to preBӧtC neurons (light grey). **B-D**) Spike rasters (each row is one burst cycle; 20 consecutive cycles are stacked) and average spike frequency for each excitatory (B), inhibitory (C), and unknown (D) rostral neuron at baseline and in SP. Time is normalized to the preBӧtC population burst duration, denoted by the x-axis tick marks.

**Figure 8:**
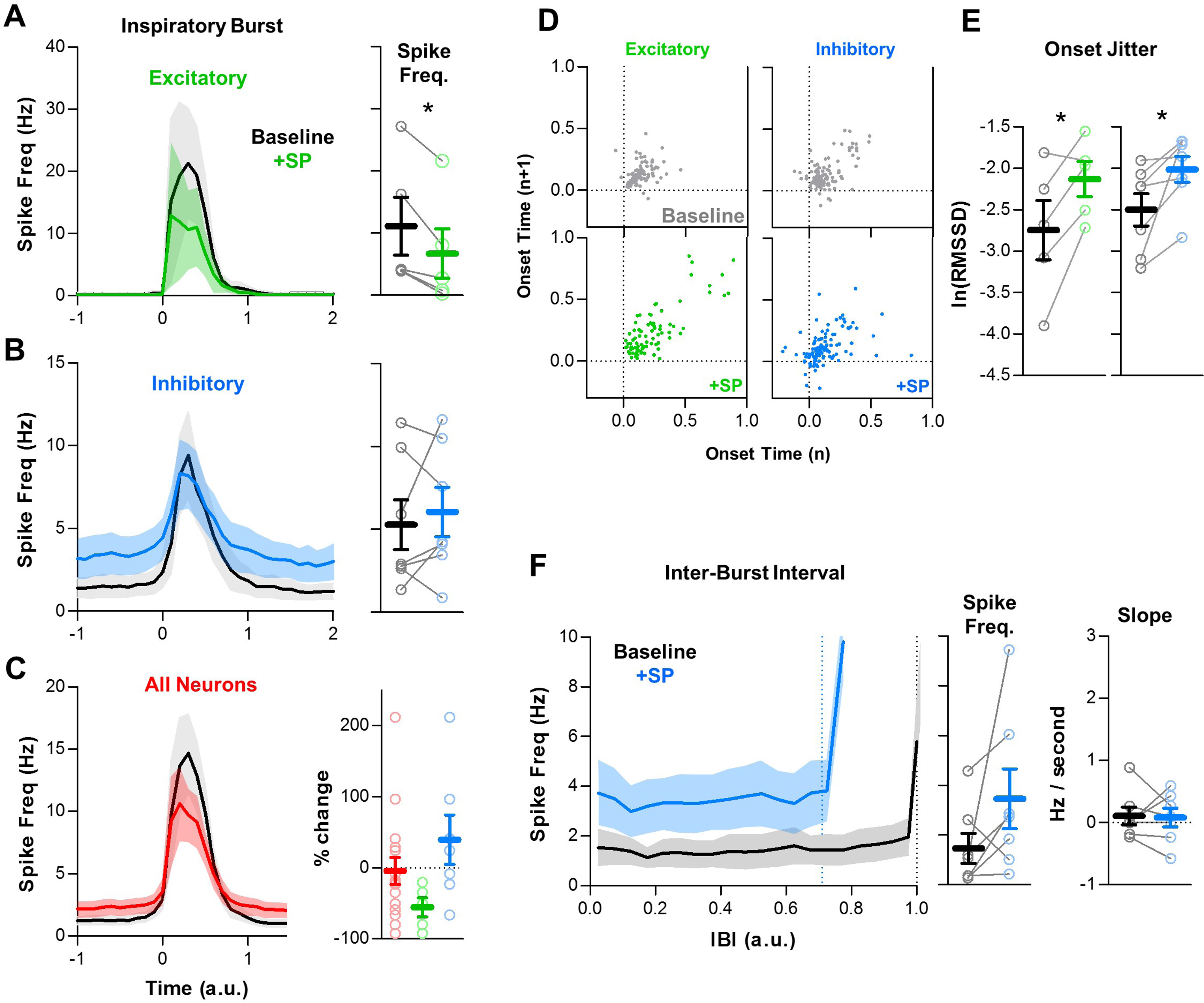
SP shifts the balance of excitation and inhibition among rostral inspiratory neurons. **A-C**) Average spike frequency as a function of time (normalized to preBӧtC population burst duration) in n=5 excitatory neurons (A), n=7 inhibitory neurons (B), and in all rostral neurons (n=16, red). Mean changes in inspiratory spike frequency at baseline and in SP are plotted to the right of each panel (paired, two tailed t-tests, A and B; one-way ANOVA, C). **D**) Poincaré plots showing burst-to-burst variability in onset times among rostral excitatory (n=5) and inhibitory (n=7) neurons (20 consecutive inspiratory bursts/neuron). **E**) Mean burst onset time variability or “jitter” at baseline and in SP for each excitatory (left) and inhibitory (right) neurons (paired t-tests). **F**) Average spike frequency of rostral inhibitory neurons during the inter-burst interval (normalized to baseline IBI) at baseline and in SP. Mean changes in spike frequency and slope during the IBI are plotted to the right (paired, two tailed t-tests).

During the inter-burst interval, the spiking activity of rostral neurons was considerably different from neurons in the preBӧtC. Unlike the pre-I spiking described for excitatory neurons in the preBӧtC (see Figs. 3 and 4), excitatory rostral neurons were generally silent during the IBI under baseline conditions, and they remained silent following application of SP (Fig. 7B). In contrast, 4 out of 7 inhibitory rostral neurons exhibited spiking during the IBI under baseline conditions, and in 3 of these neurons spike frequency during the IBI was increased by SP. Among the 3 inhibitory rostral neurons that were silent during the IBI under baseline conditions, 2 were recruited to spike during the IBI in response to SP (Overall, 6 of 7 exhibited spiking during the IBI in SP). The spiking of these neurons did not exhibit a pre-inspiratory ramp under baseline conditions (Slope: 0.11±0.14Hz/second; p>0.05) or in the presence of SP (Slope: 0.08±0.15Hz/second; p>0.05). Thus, there was a trend toward increased tonic inhibition during the IBI (1.5±0.6 to 3.5±1.2Hz; p>0.05) in the rostral inspiratory column in response to SP (Fig. 8F).

## Discussion

The rhythm generating network that produces breathing movements must constantly adjust to changing metabolic demands and also adapt to overlapping volitional and reflexive behaviors (Feldman et al., 2013, Ramirez and Baertsch, 2018b). Unravelling mechanisms that support this dynamic control may improve our understanding of disorders of the nervous system that destabilize breathing, such as Parkinson’s disease, Rett syndrome, sudden infant death syndrome, congenital central hypoventilation syndrome, multiple-systems atrophy, and amyotrophic lateral sclerosis (Oliveira et al., 2019, Ramirez et al., 2018, Schwarzacher et al., 2011, Katz et al., 2009, Moreira et al., 2016). Changes in neuromodulatory systems within the brainstem have been linked to many of these and other respiratory control disorders (Doi and Ramirez, 2008, Viemari et al., 2005). Here, we introduce the concept that neuromodulation can differentially control distinct phases of the rhythmogenic process to regulate the frequency and stability of breathing (Fig. 9).

**Figure 9:**
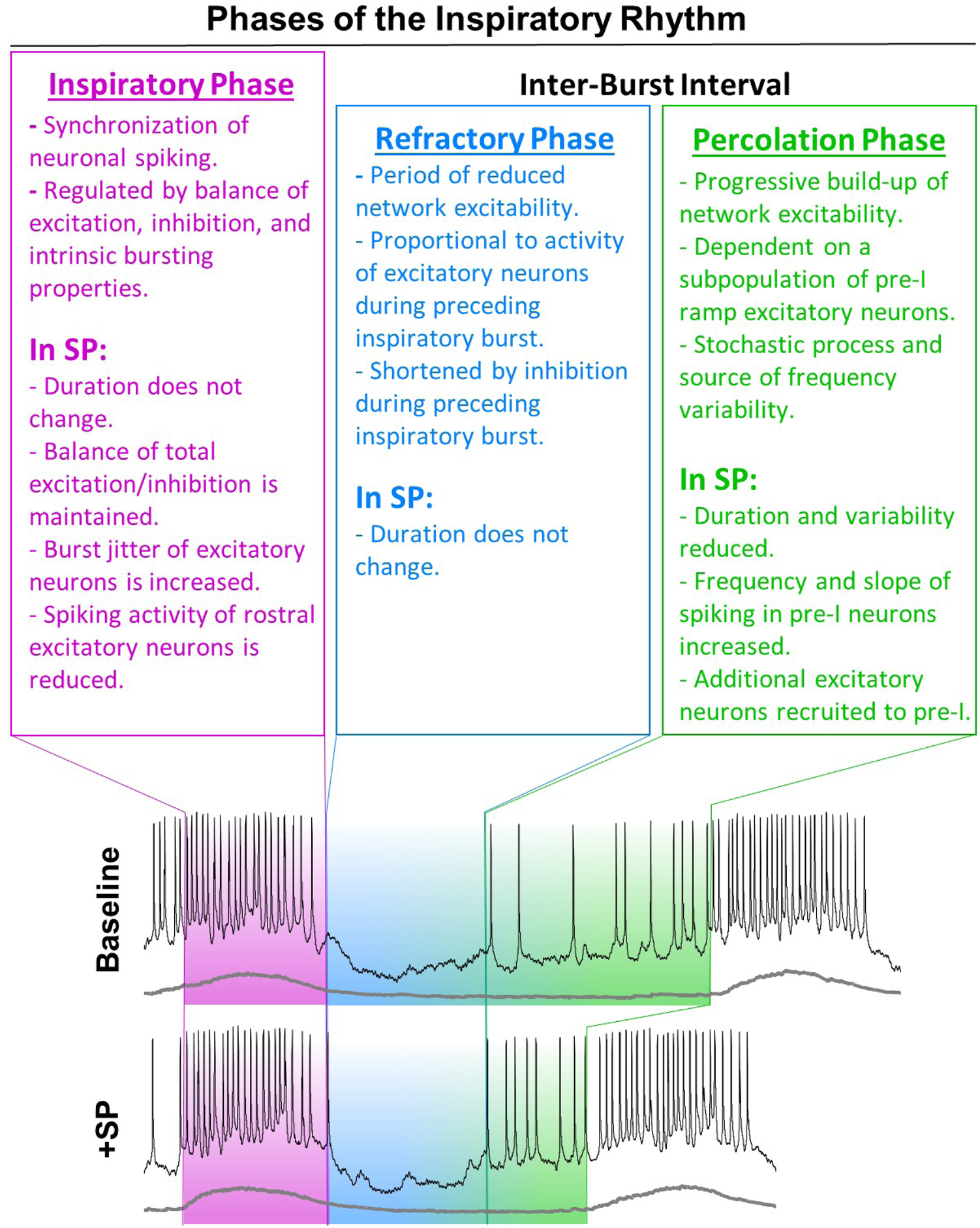
Summary schematic of inspiratory phases and their differential regulation by SP.

During the inspiratory phase, each burst is assembled stochastically via heterogeneous interactions among a combination of intertwined synaptic and intrinsic properties (Ramirez and Baertsch, 2018b). Although exclusively excitatory synaptic interactions (Kam et al., 2013a) or intrinsic bursting mechanisms (Pena et al., 2004) may be able to produce rhythm in isolation, this is unlikely to occur under normal conditions since these properties interact strongly. To the contrary, the combination of excitatory and inhibitory synaptic interactions with intrinsic bursting properties, known as the “rhythmogenic triangle” (Ramirez and Baertsch, 2018b), is critical for the flexibility of this dynamic network (Ramirez et al., 2004, Rubin and Smith, 2019). Neuromodulators play important roles in regulating both synaptic and intrinsic bursting properties, perhaps best demonstrated in invertebrate model systems. In these networks, neuromodulators can inhibit or strengthen synaptic interactions (Harris-Warrick et al., 1998, Marder et al., 2014, Nusbaum et al., 2001), as well as induce or suppress intrinsic bursting properties (Elson and Selverston, 1992, Flamm and Harris-Warrick, 1986, Zhang and Harris-Warrick, 1994). These important principles also apply to the aminergic and peptidergic modulation of the mammalian preBötC network (Doi and Ramirez, 2010). NK_1_, 5-HT2A and α2 adrenergic receptor activation modulates voltage-dependent (I_NaP_), but not voltage-independent (I_CAN_), bursting conductances, whereas α1 adrenergic receptors modulate I_CAN_-, but not I_Nap_-, dependent bursting (Pena and Ramirez, 2004, Tryba et al., 2008, Viemari and Ramirez, 2006, Pena and Ramirez, 2002). Rhythmogenesis is also modulated by sensory feedback, as demonstrated in numerous rhythmogenic networks (Ache et al., 2019, Daur et al., 2012, Grillner and El Manira, 2015, Knafo and Wyart, 2018, Vidal-Gadea et al., 2010). In the respiratory network, mechanisms of sensory feedback inhibition, such as the Breuer-Hering reflex, can increase synaptic inhibition during the inspiratory phase, which drives breathing frequency through modulation of the refractory period (Baertsch et al., 2018).

NK_1_R is expressed on preBӧtC neurons important for rhythmogenesis (Gray et al., 2001), and some evidence suggests NK_1_R is primarily expressed on excitatory neurons (Gray et al., 1999). However, it is unclear how this modulator excites the respiratory network. If SP activates excitatory neurons during inspiratory bursts, one would expect a prolongation of the refractory phase, which would limit rather than promote a frequency increase. However, we found that SP does not affect the spiking frequency of either excitatory or inhibitory preBӧtC neurons during the inspiratory phase (Figs. 3,4,5). Thus, an interesting question is: How is the balance of excitation and inhibition maintained during the inspiratory phase despite the effects of SP on excitatory preBӧtC neurons? We found that SP increases burst onset jitter specifically of excitatory neurons that lack pre-I activity (Fig. 6). This increase in timing variability implies reduced synchronization of this excitatory population during inspiratory bursts, which may be one mechanism that prevents excitatory neurons from becoming hyperactive during the inspiratory phase in response to SP.

Furthermore, recent evidence suggests that inspiratory neurons located rostral to the preBӧtC also contribute to the dynamic regulation of breathing frequency (Baertsch et al., 2019). However, under normal conditions, inhibition restrains the rhythm generating ability of these rostral neurons, as recruitment of these neurons is associated with increased excitation during inspiratory bursts, a prolonged refractory phase, and consequently a decreased respiratory frequency (Baertsch et al., 2019). Here, we found that SP has relatively heterogeneous effects on the activity of rostral inspiratory neurons; e.g. some increase, some decrease, and some do not change (Fig. 7). However, changes are more consistent specifically among rostral excitatory neurons, which exhibit decreased activity during inspiratory bursts (Fig. 8). Thus, reduced excitatory inputs from rostral neurons to the preBӧtC may be an additional mechanism that maintains the balance of excitation and inhibition during the inspiratory phase, despite SP-induced excitation. However, other mechanisms, such as depletion of excitatory synaptic vesicles (Rubin et al., 2009), may also contribute. The net effect of these processes is that the balance of excitation and inhibition during the inspiratory phase is maintained in SP.

The inspiratory phase is followed by a period of reduced excitability in the preBötC network. This refractory period is thought to arise from a combination of presynaptic depression (Kottick and Del Negro, 2015) and activation of slow hyperpolarizing current(s) (Baertsch et al., 2018, Krey et al., 2010) in glutamatergic Dbx1 neurons during inspiratory bursts. Indeed, refractoriness is maximal immediately following the inspiratory burst followed by a gradual recovery of excitability (Fig. 1), likely as vesicles are recycled and hyperpolarizing conductances are inactivated. This refractory phase manifests experimentally as a period during which the probability of evoking an ectopic inspiratory burst via optogenetic stimulation of Dbx1 neurons is reduced (Fig. 1A). However, the refractory period is not absolute as it can be overcome if the stimulus is of sufficient strength (Vann et al., 2018). Using a stimulus procedure consistent with previous reports (Baertsch et al., 2018, Baertsch et al., 2019, Kottick and Del Negro, 2015), we found that SP does not change the duration of the refractory period. This finding is consistent with our demonstration that excitatory neurons do not show an increased activation during the inspiratory phase (Figs. 3B,4B). Importantly, the minimum duration of spontaneous inter-burst intervals continues to be restrained by the refractory period in SP (Fig. 1). Thus, refractory mechanisms remain an important determinant of breathing frequency in the presence of this neuromodulator. This may be functionally important to prevent excitatory neuromodulators such as SP from driving the respiratory network out of its physiological frequency range. Indeed, neuronal networks must not only be capable of dynamically regulating their frequency, but they must also be able to maintain stability in spite of heterogeneous, intrinsically variable cellular components that receive converging inputs from numerous excitatory neuromodulators (Marder et al., 2014).

As the inspiratory network transitions out of the refractory phase, recurrent synaptic excitation, particularly involving Dbx1 neurons (Wang et al., 2014), is thought to constitute a key rhythmogenic mechanism within the preBӧtC (Del Negro et al., 2018). This process, sometimes referred to as the “group pacemaker” hypothesis (Del Negro and Hayes, 2008), involves the stochastic percolation of excitatory synaptic interactions that gradually builds-up excitability within the network between inspiratory bursts. A pre-inspiratory “ramp” in spiking activity is observed in some preBӧtC neurons as a result. We found that the magnitude and slope of this pre-inspiratory ramp, as well as the number of excitatory neurons that have pre-inspiratory activity, is increased by SP. These recruited excitatory neurons also exhibit a ramping of spiking activity during the IBI, suggesting that they participate in this potential rhythm generating network-based mechanism. Together, these effects likely increase the rate of recurrent excitation during this percolation phase (Fig. 9), which underlies the increase in breathing frequency induced by SP. Moreover, we found that, on a cycle-to-cycle basis, the slope of pre-inspiratory ramp activity is inversely related to the inter-burst interval. Thus, variations in the stochastic percolation of excitation during this phase seem to predict variability in the duration between inspiratory bursts. Our data suggest that, by increasing the rate of recurrent excitation, this process becomes more consistent and breathing irregularity is reduced. We conclude that the dual effects of SP on breathing frequency and stability are primarily a consequence of its effects on the percolation phase of the inspiratory rhythm.

Based on our collective results, we propose a conceptual framework for inspiratory rhythm generation in which three distinct phases, the inspiratory phase, refractory phase, and percolation phase, can be differentially modulated to influence breathing dynamics and stability (Fig. 9). This concept may provide a foundation for understanding breathing in the context of many other physiological and pathological conditions (Bright et al., 2018, Saito et al., 2001), and it may also serve as a guide for understanding the dynamic control rhythm generating networks in general.

## Materials and Methods

### Animals

All experiments and animal procedures were approved by the Seattle Children’s Research Institute’s Animal Care and Use Committee and conducted in accordance with the National Institutes of Health guidelines. Experiments were performed on neonatal (p6-p12) male and female C57-Bl6 mice bred at Seattle Children’s Research Institute. All mice were group housed with access to food and water *ad libitum* in a temperature controlled (22±1°C) facility with a 12hr light/dark cycle. For optogenetic experiments, *Vglut2^Cre^* and *Vgat^Cre^* (Vong et al., 2011) homozygous breeder lines were obtained from Jackson Laboratories (Stock numbers 028863 and 016962, respectively). Heterozygous *Dbx1^CreERT2^* mice were donated by Dr. Del Negro (College of William and Mary, VA) and a homozygous breeder line was generated at SCRI. *Dbx1^CreERT2^* dams were plug checked and injected at E9.5 with tamoxifen (24mg/kg, i.p.) to target preBötC neurons (Kottick et al., 2017). Cre mice were crossed with homozygous mice containing a floxed STOP channelrhodopsin2 fused to an EYFP (Ai32) reporter sequence (JAX #024109). Male and female offspring were chosen at random based on litter distributions.

### In-vitro medullary horizontal slice preparations

Horizontal medullary slices were prepared from postnatal day 6-12 mice as described in detail previously (Anderson et al., 2016; Baertsch et al., 2019). Brainstems were dissected in ice cold artificial cerebrospinal fluid (aCSF; in mM: 118 NaCl, 3.0 KCl, 25 NaHCO_3_, 1 NaH_2_PO_4_, 1.0 MgCl_2_, 1.5 CaCl_2_, 30 D-glucose) equilibrated with carbogen (95% O_2_, 5% CO_2_). When equilibrated with gas mixtures containing 5% CO_2_ at ambient pressure, aCSF had an osmolarity of 305–312mOSM and a pH of 7.40– 7.45. The dorsal surface of each brainstem was secured with super glue to an agar block cut at a ∼15° angle (rostral end facing up). Brainstems were first sectioned in the transverse plane (200µm steps) using a vibratome (Leica 1000S) until the VII nerves were visualized. The agar block was then reoriented to position the ventral surface of the brainstem facing up with the rostral end towards the vibratome blade to section the brainstem in the horizontal plane. The blade was leveled with the ventral edge of the brainstem and a single ∼850µm step was taken to create the horizontal slice.

Slices were placed in a custom recording chamber containing circulating aCSF (∼15ml/min) warmed to 30°C. The [K+] in the aCSF was then gradually raised from 3mM to 8mM over ∼10min to boost neuronal excitability. Rhythmic extracellular neuronal population activity was recorded by positioning polished glass pipettes (<1MΩ tip resistance) filled with aCSF on the surface of the slice. Signals were amplified 10,000X, filtered (low pass, 300Hz; high pass, 5kHz), rectified, integrated, and digitized (Digidata 1550A, Axon Instruments). The activity of single neurons was recorded using the blind patch clamp approach. Recording electrodes were pulled from borosilicate glass (4-8MΩ tip resistance) using a P-97 Flaming/Brown micropipette puller (Sutter Instrument Co., Novato, CA) and filled with intracellular patch electrode solution containing (in mM): 140 potassium gluconate, 1 CaCl_2_, 10 EGTA, 2 MgCl_2_, 4 Na_2_ATP, and 10 Hepes (pH 7.2). To map the location of recorded neurons, patch pipettes were backfilled with intracellular patch solution containing 2mg/ml Alexa Fluor568 Hyrdrazide (ThermoFisher). Neuronal spiking activity was recorded in whole-cell or cell-attached configuration with a multiclamp amplifier in current clamp mode (Molecular Devices, Sunnyvale, CA). Extracellular and intracellular signals were acquired in pCLAMP software (Molecular Devices, Sunnyvale, CA). Immediately following electrophysiology experiments, fresh, unfixed slices were imaged to determine the location(s) of the intracellular recording sites.

### Optogenetic and pharmacological manipulations

A 200µm diameter glass fiber optic (0.24NA) connected to a blue (470nm) high-powered LED was positioned above the preBötC contralateral to the extracellular electrode and ipsilateral to the intracellular electrode. Power was set ≤1mW/mm^2^. To determine the probability of light-evoking inspiratory bursts, 200ms light pulses were TTL-triggered every 20s to stimulate Dbx1 neurons (≥50 trails per experiment). Trials were excluded from the analysis if the light pulse occurred during an ongoing spontaneous inspiratory burst. During most intracellular recordings, neurons were classified as excitatory or inhibitory using an optogenetic approach. In Vgat^Cre^;Rosa26^ChR2-EYFP^ slices, neurons that depolarized during photostimulation were classified as inhibitory, while those that hyperpolarized or did not respond were presumed to be excitatory. Because a depolarizing response to stimulation of excitatory neurons could be driven synaptically instead of from channelrhodopsin2 expression directly, in Vglut2^Cre^;Rosa26^ChR2-EYFP^ and Dbx1^CreERT2^;Rosa26^ChR2-EYFP^ slices, neurons were classified as excitatory or inhibitory based on the presence or absence of a depolarizing response to light, respectively, following pharmacological blockade of excitatory AMPAR- and NMDAR-dependent synaptic transmission (20µM CNQX, 20µM CPP).

Substance P was purchased from Tocris (Cat#: 1156), diluted in water to a concentration of 5mM, and stored in stock aliquots at -20°C. In all experiments, a ∼10min baseline period of stable inspiratory activity was recorded prior to bath application of substance P to 0.5-1.0µM. Intracellular and extracellular population activity was then recorded for >10min prior to washout into fresh aCSF.

### Microscopy

2.5X brightfield and epifluorescent images of the dorsal surface of horizontal slices were acquired on a Leica DM 4000 B epifluorescence microscope. Following intracellular recording experiments, the location of each recorded neuron within the horizontal slice was immediately quantified by overlaying the brightfield and an epifluorescent image of Alexa Fluor 568 labelled cell(s). Images were then traced in powerpoint and overlaid with the midline and rostral edge (VII nerve) aligned to show the relative locations of recorded cells (see Figs 2A and 7A).

### Statistical analysis

Data was analyzed using Clampfit software (Molecular Devices). Integrated population bursts and individual action potentials were detected using Clampfit’s peak-detection analysis. Statistical analyses were performed using GraphPad Prism6 software and are detailed for each experiment in the Figure Legends. Groups were compared using appropriate two-tailed t-tests, or one-way ANOVAs with Bonferonni’s multiple comparisons post hoc tests. Welch’s correction was used for unequal variances where appropriate. Non-linear regression analysis was used to determine differences between probability curves (Fig 1). Linear regression analyses were used to determine relationships between inter-burst intervals and pre-inspiratory ramp slope (Figs 3G and 5F). Differences were considered significant at p<0.05 and data are displayed as individual data points with overlaid means±SE. Significance is denoted in the figures as follows: **** p<0.0001; *** p<0.001; ** p<0.01; * p<0.05. Experimenters were not blinded during data collection or analysis.

## Acknowledgments

We thank NIH grants R01 HL126523 (Awarded to JMR), R01 HL144801 (Awarded to JMR), P01 HL 090554 (Awarded to JMR), K99 HL145004 (Awarded to NAB) and F32 HL134207 (Awarded to NAB) for funding this project.

## Author Contributions

Conceptualization, N.A.B. and J.M.R.; Methodology, N.A.B., Investigation, N.A.B., Formal Analysis, N.A.B.; Writing–Original Draft, N.A.B.; Writing–Review & Editing, N.A.B., and J.M.R.; Funding Acquisition, N.A.B. and J.M.R.; Visualization, N.A.B.; Supervision, J.M.R.

## References

Ache, J. M., Namiki, S., Lee, A., Branson, K. & Card, G. M. 2019. State-dependent decoupling of sensory and motor circuits underlies behavioral flexibility in Drosophila. Nat Neurosci, 22, 1132–1139.

Anderson, T. M., Garcia, A. J., 3rd, Baertsch, N. A., Pollak, J., Bloom, J. C., Wei, A. D., Rai, K. G. & Ramirez, J. M. 2016. A novel excitatory network for the control of breathing. Nature, 536, 76–80.

Baertsch, N. A., Baertsch, H. C. & Ramirez, J. M. 2018. The interdependence of excitation and inhibition for the control of dynamic breathing rhythms. Nat Commun, 9, 843.

Baertsch, N. A., Severs, L. J., Anderson, T. M. & Ramirez, J. M. 2019. A spatially dynamic network underlies the generation of inspiratory behaviors. Proc Natl Acad Sci U S A, 116, 7493–7502.

Barnes, B. J., Tuong, C. M. & Mellen, N. M. 2007. Functional imaging reveals respiratory network activity during hypoxic and opioid challenge in the neonate rat tilted sagittal slab preparation. J Neurophysiol, 97, 2283–92.

Basar, E. & Duzgun, A. 2016. Links of Consciousness, Perception, and Memory by Means of Delta Oscillations of Brain. Front Psychol, 7, 275.

Basar, E. & Guntekin, B. 2008. A review of brain oscillations in cognitive disorders and the role of neurotransmitters. Brain Res, 1235, 172–93.

Ben-Mabrouk, F. & Tryba, A. K. 2010. Substance P modulation of TRPC3/7 channels improves respiratory rhythm regularity and ICAN-dependent pacemaker activity. Eur J Neurosci, 31, 1219–32.

Bouvier, J., Thoby-Brisson, M., Renier, N., Dubreuil, V., Ericson, J., Champagnat, J., Pierani, A., Chedotal, A. & Fortin, G. 2010. Hindbrain interneurons and axon guidance signaling critical for breathing. Nat Neurosci, 13, 1066–74.

Bright, F. M., Vink, R. & Byard, R. W. 2018. The potential role of substance P in brainstem homeostatic control in the pathogenesis of sudden infant death syndrome (SIDS). Neuropeptides, 70, 1–8.

Brittain, J. S., Sharott, A. & Brown, P. 2014. The highs and lows of beta activity in cortico-basal ganglia loops. Eur J Neurosci, 39, 1951–9.

Carroll, M. S. & Ramirez, J. M. 2013. Cycle-by-cycle assembly of respiratory network activity is dynamic and stochastic. J Neurophysiol, 109, 296–305.

Carroll, M. S., Viemari, J. C. & Ramirez, J. M. 2013. Patterns of inspiratory phase-dependent activity in the in vitro respiratory network. J Neurophysiol, 109, 285–95.

Chen, Z., Hedner, J. & Hedner, T. 1990. Substance P in the ventrolateral medulla oblongata regulates ventilatory responses. J Appl Physiol (1985), 68, 2631–9.

Cohen, M. I. 1981. Central determinants of respiratory rhythm. Annu Rev Physiol, 43, 91–104.

Colgin, L. L. 2016. Rhythms of the hippocampal network. Nat Rev Neurosci, 17, 239–49.

Daur, N., Diehl, F., Mader, W. & Stein, W. 2012. The stomatogastric nervous system as a model for studying sensorimotor interactions in real-time closed-loop conditions. Front Comput Neurosci, 6, 13.

Del Negro, C. A., Funk, G. D. & Feldman, J. L. 2018. Breathing matters. Nat Rev Neurosci, 19, 351–367.

Del Negro, C. A. & Hayes, J. A. 2008. A ‘group pacemaker’ mechanism for respiratory rhythm generation. J Physiol, 586, 2245–6.

Del Negro, C. A., Hayes, J. A., Pace, R. W., Brush, B. R., Teruyama, R. & Feldman, J. L. 2010. Synaptically activated burst-generating conductances may underlie a group-pacemaker mechanism for respiratory rhythm generation in mammals. Prog Brain Res, 187, 111–36.

Dick, T. E., Dutschmann, M., Feldman, J. L., Fong, A. Y., Hulsmann, S., Morris, K. M., Ramirez, J. M. & Smith, J. C. 2018. Facts and challenges in respiratory neurobiology. Respir Physiol Neurobiol, 258, 104–107.

Doi, A. & Ramirez, J. M. 2008. Neuromodulation and the orchestration of the respiratory rhythm. Respir Physiol Neurobiol, 164, 96–104.

Doi, A. & Ramirez, J. M. 2010. State-dependent interactions between excitatory neuromodulators in the neuronal control of breathing. J Neurosci, 30, 8251–62.

Elson, R. C. & Selverston, A. I. 1992. Mechanisms of gastric rhythm generation in the isolated stomatogastric ganglion of spiny lobsters: bursting pacemaker potentials, synaptic interactions, and muscarinic modulation. J Neurophysiol, 68, 890–907.

Ezure, K. 1990. Synaptic connections between medullary respiratory neurons and considerations on the genesis of respiratory rhythm. Prog Neurobiol, 35, 429–50.

Feldman, J. L., Del Negro, C. A. & Gray, P. A. 2013. Understanding the rhythm of breathing: so near, yet so far. Annu Rev Physiol, 75, 423–52.

Feldman, J. L. & Kam, K. 2015. Facing the challenge of mammalian neural microcircuits: taking a few breaths may help. J Physiol, 593, 3–23.

Flamm, R. E. & Harris-Warrick, R. M. 1986. Aminergic modulation in lobster stomatogastric ganglion. II. Target neurons of dopamine, octopamine, and serotonin within the pyloric circuit. J Neurophysiol, 55, 866–81.

Ge, Q. & Feldman, J. L. 1998. AMPA receptor activation and phosphatase inhibition affect neonatal rat respiratory rhythm generation. J Physiol, 509 (Pt 1), 255–66.

Golombek, D. A., Bussi, I. L. & Agostino, P. V. 2014. Minutes, days and years: molecular interactions among different scales of biological timing. Philos Trans R Soc Lond B Biol Sci, 369, 20120465.

Gray, P. A., Hayes, J. A., Ling, G. Y., Llona, I., Tupal, S., Picardo, M. C., Ross, S. E., Hirata, T., Corbin, J. G., Eugenin, J. & Del Negro, C. A. 2010. Developmental origin of preBotzinger complex respiratory neurons. J Neurosci, 30, 14883–95.

Gray, P. A., Janczewski, W. A., Mellen, N., Mccrimmon, D. R. & Feldman, J. L. 2001. Normal breathing requires preBotzinger complex neurokinin-1 receptor-expressing neurons. Nat Neurosci, 4, 927–30.

Gray, P. A., Rekling, J. C., Bocchiaro, C. M. & Feldman, J. L. 1999. Modulation of respiratory frequency by peptidergic input to rhythmogenic neurons in the preBotzinger complex. Science, 286, 1566–8.

Grillner, S. & El Manira, A. 2015. The intrinsic operation of the networks that make us locomote. Curr Opin Neurobiol, 31, 244–9.

Hanslmayr, S., Staresina, B. P. & Bowman, H. 2016. Oscillations and Episodic Memory: Addressing the Synchronization/Desynchronization Conundrum. Trends Neurosci, 39, 16–25.

Harris-Warrick, R. M., Johnson, B. R., Peck, J. H., Kloppenburg, P., Ayali, A. & Skarbinski, J. 1998. Distributed effects of dopamine modulation in the crustacean pyloric network. Ann N Y Acad Sci, 860, 155–67.

Harris, K. D., Dashevskiy, T., Mendoza, J., Garcia, A. J., 3rd, Ramirez, J. M. & Shea-Brown, E. 2017. Different roles for inhibition in the rhythm-generating respiratory network. J Neurophysiol, 118, 2070–2088.

Hayes, J. A. & Del Negro, C. A. 2007. Neurokinin receptor-expressing pre-botzinger complex neurons in neonatal mice studied in vitro. J Neurophysiol, 97, 4215–24.

Jenkin, S. E. & Milsom, W. K. 2014. Expiration: breathing’s other face. Prog Brain Res, 212, 131–47.

Kam, K., Worrell, J. W., Janczewski, W. A., Cui, Y. & Feldman, J. L. 2013a. Distinct inspiratory rhythm and pattern generating mechanisms in the preBotzinger complex. J Neurosci, 33, 9235–45.

Kam, K., Worrell, J. W., Ventalon, C., Emiliani, V. & Feldman, J. L. 2013b. Emergence of population bursts from simultaneous activation of small subsets of preBotzinger complex inspiratory neurons. J Neurosci, 33, 3332–8.

Katz, D. M., Dutschmann, M., Ramirez, J. M. & Hilaire, G. 2009. Breathing disorders in Rett syndrome: progressive neurochemical dysfunction in the respiratory network after birth. Respir Physiol Neurobiol, 168, 101–8.

Kiehn, O. 2016. Decoding the organization of spinal circuits that control locomotion. Nat Rev Neurosci, 17, 224–38.

Knafo, S. & Wyart, C. 2018. Active mechanosensory feedback during locomotion in the zebrafish spinal cord. Curr Opin Neurobiol, 52, 48–53.

Kottick, A. & Del Negro, C. A. 2015. Synaptic Depression Influences Inspiratory-Expiratory Phase Transition in Dbx1 Interneurons of the preBotzinger Complex in Neonatal Mice. J Neurosci, 35, 11606–11.

Kottick, A., Martin, C. A. & Del Negro, C. A. 2017. Fate mapping neurons and glia derived from Dbx1-expressing progenitors in mouse preBotzinger complex. Physiol Rep, 5.

Krey, R. A., Goodreau, A. M., Arnold, T. B. & Del Negro, C. A. 2010. Outward Currents Contributing to Inspiratory Burst Termination in preBotzinger Complex Neurons of Neonatal Mice Studied in Vitro. Front Neural Circuits, 4, 124.

Long, S. & Duffin, J. 1986. The neuronal determinants of respiratory rhythm. Prog Neurobiol, 27, 101–82.

Mantyh, P. W. 2002. Neurobiology of substance P and the NK1 receptor. J Clin Psychiatry, 63 Suppl 11, 6–10.

Marder, E., O’leary, T. & Shruti, S. 2014. Neuromodulation of circuits with variable parameters: single neurons and small circuits reveal principles of state-dependent and robust neuromodulation. Annu Rev Neurosci, 37, 329–46.

Milsom, W. K. 1991. Intermittent breathing in vertebrates. Annu Rev Physiol, 53, 87–105.

Moore, J. D., Deschenes, M., Furuta, T., Huber, D., Smear, M. C., Demers, M. & Kleinfeld, D. 2013. Hierarchy of orofacial rhythms revealed through whisking and breathing. Nature, 497, 205–10.

Moreira, T. S., Takakura, A. C., Czeisler, C. & Otero, J. J. 2016. Respiratory and autonomic dysfunction in congenital central hypoventilation syndrome. J Neurophysiol, 116, 742–52.

Nakamura, Y., Katakura, N., Nakajima, M. & Liu, J. 2004. Rhythm generation for food-ingestive movements. Prog Brain Res, 143, 97–103.

Narayanan, N. S. & Dileone, R. J. 2017. Lip Sync: Gamma Rhythms Orchestrate Top-Down Control of Feeding Circuits. Cell Metab, 25, 497–498.

Neske, G. T. 2015. The Slow Oscillation in Cortical and Thalamic Networks: Mechanisms and Functions. Front Neural Circuits, 9, 88.

Nusbaum, M. P., Blitz, D. M., Swensen, A. M., Wood, D. & Marder, E. 2001. The roles of co-transmission in neural network modulation. Trends Neurosci, 24, 146–54.

Oke, Y., Miwakeichi, F., Oku, Y., Hirrlinger, J. & Hulsmann, S. 2018. Cell Type-Dependent Activation Sequence During Rhythmic Bursting in the PreBotzinger Complex in Respiratory Rhythmic Slices From Mice. Front Physiol, 9, 1219.

Oliveira, L. M., Oliveira, M. A., Moriya, H. T., Moreira, T. S. & Takakura, A. C. 2019. Respiratory disturbances in a mouse model of Parkinson’s disease. Exp Physiol, 104, 729–739.

Palva, S. & Palva, J. M. 2018. Roles of Brain Criticality and Multiscale Oscillations in Temporal Predictions for Sensorimotor Processing. Trends Neurosci, 41, 729–743.

Paton, J. J. & Buonomano, D. V. 2018. The Neural Basis of Timing: Distributed Mechanisms for Diverse Functions. Neuron, 98, 687–705.

Pena, F., Parkis, M. A., Tryba, A. K. & Ramirez, J. M. 2004. Differential contribution of pacemaker properties to the generation of respiratory rhythms during normoxia and hypoxia. Neuron, 43, 105–17.

Pena, F. & Ramirez, J. M. 2002. Endogenous activation of serotonin-2A receptors is required for respiratory rhythm generation in vitro. J Neurosci, 22, 11055–64.

Pena, F. & Ramirez, J. M. 2004. Substance P-mediated modulation of pacemaker properties in the mammalian respiratory network. J Neurosci, 24, 7549–56.

Picardo, M. C., Weragalaarachchi, K. T., Akins, V. T. & Del Negro, C. A. 2013. Physiological and morphological properties of Dbx1-derived respiratory neurons in the pre-Botzinger complex of neonatal mice. J Physiol, 591, 2687–703.

Ptak, K., Burnet, H., Blanchi, B., Sieweke, M., De Felipe, C., Hunt, S. P., Monteau, R. & Hilaire, G. 2002. The murine neurokinin NK1 receptor gene contributes to the adult hypoxic facilitation of ventilation. Eur J Neurosci, 16, 2245–52.

Ptak, K., Yamanishi, T., Aungst, J., Milescu, L. S., Zhang, R., Richerson, G. B. & Smith, J. C. 2009. Raphe neurons stimulate respiratory circuit activity by multiple mechanisms via endogenously released serotonin and substance P. J Neurosci, 29, 3720–37.

Ramirez, J. M. & Baertsch, N. 2018a. Defining the Rhythmogenic Elements of Mammalian Breathing. Physiology (Bethesda), 33, 302–316.

Ramirez, J. M. & Baertsch, N. A. 2018b. The Dynamic Basis of Respiratory Rhythm Generation: One Breath at a Time. Annu Rev Neurosci, 41, 475–499.

Ramirez, J. M., Dashevskiy, T., Marlin, I. A. & Baertsch, N. 2016. Microcircuits in respiratory rhythm generation: commonalities with other rhythm generating networks and evolutionary perspectives. Curr Opin Neurobiol, 41, 53–61.

Ramirez, J. M., Ramirez, S. C. & Anderson, T. M. 2018. Sudden Infant Death Syndrome, Sleep, and the Physiology and Pathophysiology of the Respiratory Network. In: Duncan, J. R. & Byard, R. W. (eds.) SIDS Sudden Infant and Early Childhood Death: The Past, the Present and the Future. Adelaide (AU): University of Adelaide Press

Ramirez, J. M., Ramirez, S. C. & Anderson, T. M. 2018 The Contributors, with the exception of which is by Federal United States employees and is therefore in the public domain.

Ramirez, J. M., Tryba, A. K. & Pena, F. 2004. Pacemaker neurons and neuronal networks: an integrative view. Curr Opin Neurobiol, 14, 665–74.

Rubin, J. E., Hayes, J. A., Mendenhall, J. L. & Del Negro, C. A. 2009. Calcium-activated nonspecific cation current and synaptic depression promote network-dependent burst oscillations. Proc Natl Acad Sci U S A, 106, 2939–44.

Rubin, J. E. & Smith, J. C. 2019. Robustness of respiratory rhythm generation across dynamic regimes. PLoS Comput Biol, 15, e1006860.

Saito, Y., Ito, M., Ozawa, Y., Matsuishi, T., Hamano, K. & Takashima, S. 2001. Reduced expression of neuropeptides can be related to respiratory disturbances in Rett syndrome. Brain Dev, 23 Suppl 1, S122–6.

Schwab, D. J., Bruinsma, R. F., Feldman, J. L. & Levine, A. J. 2010. Rhythmogenic neuronal networks, emergent leaders, and k-cores. Phys Rev E Stat Nonlin Soft Matter Phys, 82, 051911.

Schwarzacher, S. W., Rub, U. & Deller, T. 2011. Neuroanatomical characteristics of the human pre-Botzinger complex and its involvement in neurodegenerative brainstem diseases. Brain, 134, 24–35.

Smith, J. C., Ellenberger, H. H., Ballanyi, K., Richter, D. W. & Feldman, J. L. 1991. Pre-Botzinger complex: a brainstem region that may generate respiratory rhythm in mammals. Science, 254, 726–9.

Tan, W., Janczewski, W. A., Yang, P., Shao, X. M., Callaway, E. M. & Feldman, J. L. 2008. Silencing preBotzinger complex somatostatin-expressing neurons induces persistent apnea in awake rat. Nat Neurosci, 11, 538–40.

Tryba, A. K., Pena, F., Lieske, S. P., Viemari, J. C., Thoby-Brisson, M. & Ramirez, J. M. 2008. Differential modulation of neural network and pacemaker activity underlying eupnea and sigh-breathing activities. J Neurophysiol, 99, 2114–25.

Vann, N. C., Pham, F. D., Dorst, K. E. & Del Negro, C. A. 2018. Dbx1 Pre-Botzinger Complex Interneurons Comprise the Core Inspiratory Oscillator for Breathing in Unanesthetized Adult Mice. eNeuro, 5.

Vidal-Gadea, A. G., Jing, X. J., Simpson, D., Dewhirst, O. P., Kondoh, Y., Allen, R. & Newland, P. L. 2010. Coding characteristics of spiking local interneurons during imposed limb movements in the locust. J Neurophysiol, 103, 603–15.

Viemari, J. C. & Ramirez, J. M. 2006. Norepinephrine differentially modulates different types of respiratory pacemaker and nonpacemaker neurons. J Neurophysiol, 95, 2070–82.

Viemari, J. C., Roux, J. C., Tryba, A. K., Saywell, V., Burnet, H., Pena, F., Zanella, S., Bevengut, M., Barthelemy-Requin, M., Herzing, L. B., Moncla, A., Mancini, J., Ramirez, J. M., Villard, L. & Hilaire, G. 2005. Mecp2 deficiency disrupts norepinephrine and respiratory systems in mice. J Neurosci, 25, 11521–30.

Vong, L., Ye, C., Yang, Z., Choi, B., Chua, S., Jr. & Lowell, B. B. 2011. Leptin action on GABAergic neurons prevents obesity and reduces inhibitory tone to POMC neurons. Neuron, 71, 142–54.

Wang, X., Hayes, J. A., Revill, A. L., Song, H., Kottick, A., Vann, N. C., Lamar, M. D., Picardo, M. C., Akins, V. T., Funk, G. D. & Del Negro, C. A. 2014. Laser ablation of Dbx1 neurons in the pre-Botzinger complex stops inspiratory rhythm and impairs output in neonatal mice. Elife, 3, e03427.

Winter, S. M., Fresemann, J., Schnell, C., Oku, Y., Hirrlinger, J. & Hulsmann, S. 2009. Glycinergic interneurons are functionally integrated into the inspiratory network of mouse medullary slices. Pflugers Arch, 458, 459–69.

Wu, J., Capelli, P., Bouvier, J., Goulding, M., Arber, S. & Fortin, G. 2017. A V0 core neuronal circuit for inspiration. Nat Commun, 8, 544.

Wyart, C. 2018. Taking a Big Step towards Understanding Locomotion. Trends Neurosci, 41, 869–870.

Wyman, R. J. 1977. Neural generation of the breathing rhythm. Annu Rev Physiol, 39, 417–48.

Yeh, S. Y., Huang, W. H., Wang, W., Ward, C. S., Chao, E. S., Wu, Z., Tang, B., Tang, J., Sun, J. J., Esther van der Heijden, M., Gray, P. A., Xue, M., Ray, R. S., Ren, D. & Zoghbi, H. Y. 2017. Respiratory Network Stability and Modulatory Response to Substance P Require Nalcn. Neuron, 94, 294–303.e4.

Zerlaut, Y. & Destexhe, A. 2017. Enhanced Responsiveness and Low-Level Awareness in Stochastic Network States. Neuron, 94, 1002–1009.

Zhang, B. & Harris-Warrick, R. M. 1994. Multiple receptors mediate the modulatory effects of serotonergic neurons in a small neural network. J Exp Biol, 190, 55–77.

Zuperku, E. J., Stucke, A. G., Hopp, F. A. & Stuth, E. A. 2017. Characteristics of breathing rate control mediated by a subregion within the pontine parabrachial complex. J Neurophysiol, 117, 1030–1042.

Zuperku, E. J., Stucke, A. G., Krolikowski, J. G., Tomlinson, J., Hopp, F. A. & Stuth, E. A. 2019. Inputs to medullary respiratory neurons from a pontine subregion that controls breathing frequency. Respir Physiol Neurobiol, 265, 127–140.

